# 4R-Tobacco-Cembranoid is a Positive Allosteric Modulator of the Alpha7 Nicotinic Acetylcholine Receptor that modulates inflammation-induced thermal hypersensitivity in mice

**DOI:** 10.1101/2022.01.22.477375

**Authors:** Luis G. Rivera-García, Adela M. Francis-Malavé, Torri D. Wilson, P. A. Ferchmin, Vesna Eterovic, Yarimar Carrasquillo

## Abstract

Alpha7 nicotinic acetylcholine receptors (α7nAChRs) are activated in response to inflammation and modulate pain in humans and rodent models. The use of α7nAChRs agonists as a therapeutic option for inflammation and pain is challenged by unwanted effects resulting from constant activation and/or desensitization of α7nAChRs. Positive allosteric modulators (PAMs) represent a compelling alternative as they increase endogenous nicotinic transmission but do not result in progressive desensitization or loss of receptor function. In the present study, we evaluated the function of the 4R tobacco cembranoid (4R) as a PAM of α7nAChR that reduces inflammation and pain-related behaviors in mouse models of inflammatory pain. Our electrophysiological experiments show that 4R potentiates choline-evoked currents in SH-SY5Y cells overexpressing α7nAChRs in a dose-dependent manner. At the behavioral level, we show that subcutaneous administration of 4R decreases inflammation-induced thermal but not tactile hypersensitivity or formalin-induced spontaneous nociceptive responses in both male and female mice. We further show reduced inflammation-induced paw edema in 4R-treated males, with no measurable effect observed in female mice. Altogether, the results from the experiments in this study identify 4R as a PAM of α7nAChRs that reduces thermal hypersensitivity in male and female mice and inflammation in a sex-specific manner. These findings highlight the use of 4R as a potential novel treatment strategy for pain and inflammation.

## Introduction

Malfunctioning or persistent activation of nociceptive systems can lead to chronic pathological pain that no longer has an adaptive function. Current options for pathological pain treatment are limited and are typically accompanied by unwanted side effects. For this reason, pain is cited as one of the main reasons for patients to seek medical attention (Schappert and Burt 2006), highlighting the need for improved treatment options.

Nicotinic acetylcholine receptors (nAChRs) are ligand-gated ion channels expressed in neuronal and non-neuronal cells across the central and peripheral nervous systems that have received attention over the years as potential targets for pain treatment (Bektas et al. 2020). The use of nAChR agonists for pain treatment, however, has been hampered by unwanted side effects that are mainly attributed to receptor desensitization or changes in expression levels as a consequence of long-term agonist administration (Quick and Lester 2002). Positive allosteric modulators (PAMs) of nAChRs that bind to a receptor at an allosteric site became a compelling alternative to synergize and augment natural cholinergic signals instead of competing against or attempting to replace them (Williams et al. 2011). PAMs are synthetic or natural molecules that exhibit pharmacological advantages over agonists. These advantages include maintaining the typical spatial and temporal pattern of endogenous neurotransmission and higher receptor subtype selectivity that result in beneficial outcomes with less unfavorable effects under clinical conditions (Uteshev 2014).

α7nAChRs are among the nicotinic receptors proposed to function as potential targets for pain treatment (Hone and McIntosh 2018, Naser and Kuner 2018, Bektas et al. 2020). Consistent with this, treatment with selective α7nAChR agonists or PAMs have been shown to provide analgesia and reduce pain-related negative affective behaviors in several inflammatory and visceral pain models (Damaj et al. 2000, Gurun et al. 2009, Rowley et al. 2010, Targowska-Duda et al. 2014, Bagdas et al. 2016). In the present study, we expand on these previous findings by identifying 4R as a novel PAM for α7nAChRs that decreases inflammation and inflammation-induced thermal hypersensitivity in mice.

Some species of Nicotiana plants, commonly referred to as tobacco plants, produce cyclic diterpenoids, called cembranoids, that interact with nAChRs (Ferchmin et al. 2009). The tobacco cembranoid used in this work is (1S, 2E, 4R, 6R, 7E, 11E)-cembra-2,7,11-triene-4,6-diol, abbreviated as 4R. It was isolated from tobacco and cigarette smoke, has the molecular formula C_20_H_34_O_2_ and the chemical structure shown in **Fig. 1a** (Saito et al. 1985). Previous studies have shown that 4R inhibits three human nAChR subtypes expressed in cell lines: neuronal α4β2 (IC_50_=19 μM) and α3β4 (IC_50_=2 μM) and the embryonic muscle AChR α1β1γδ-AChR, demonstrating that 4R is a nicotinic ligand (Ferchmin et al. 2001). Additional studies have further shown that 4R has anti-inflammatory effects and is neuroprotective against excitotoxicity in-vivo in rodents and also ex-vivo in hippocampal slices (Ferchmin et al. 2005, Eterović et al. 2011, Ferchmin et al. 2014, Ferchmin et al. 2015, Martins et al. 2015). The potential use of 4R for pain treatment, however, has not been evaluated. Here, we confirm the anti-inflammatory effects of 4R in a mouse model of inflammatory pain and further demonstrate that inflammation-induced thermal hypersensitivity is decreased with 4R via positive allosteric modulation of α7nAChRs. Together, our results identified a novel analgesic function for 4R and further support the use of nAChRs PAMs for pain treatment.

**Figure 1:**
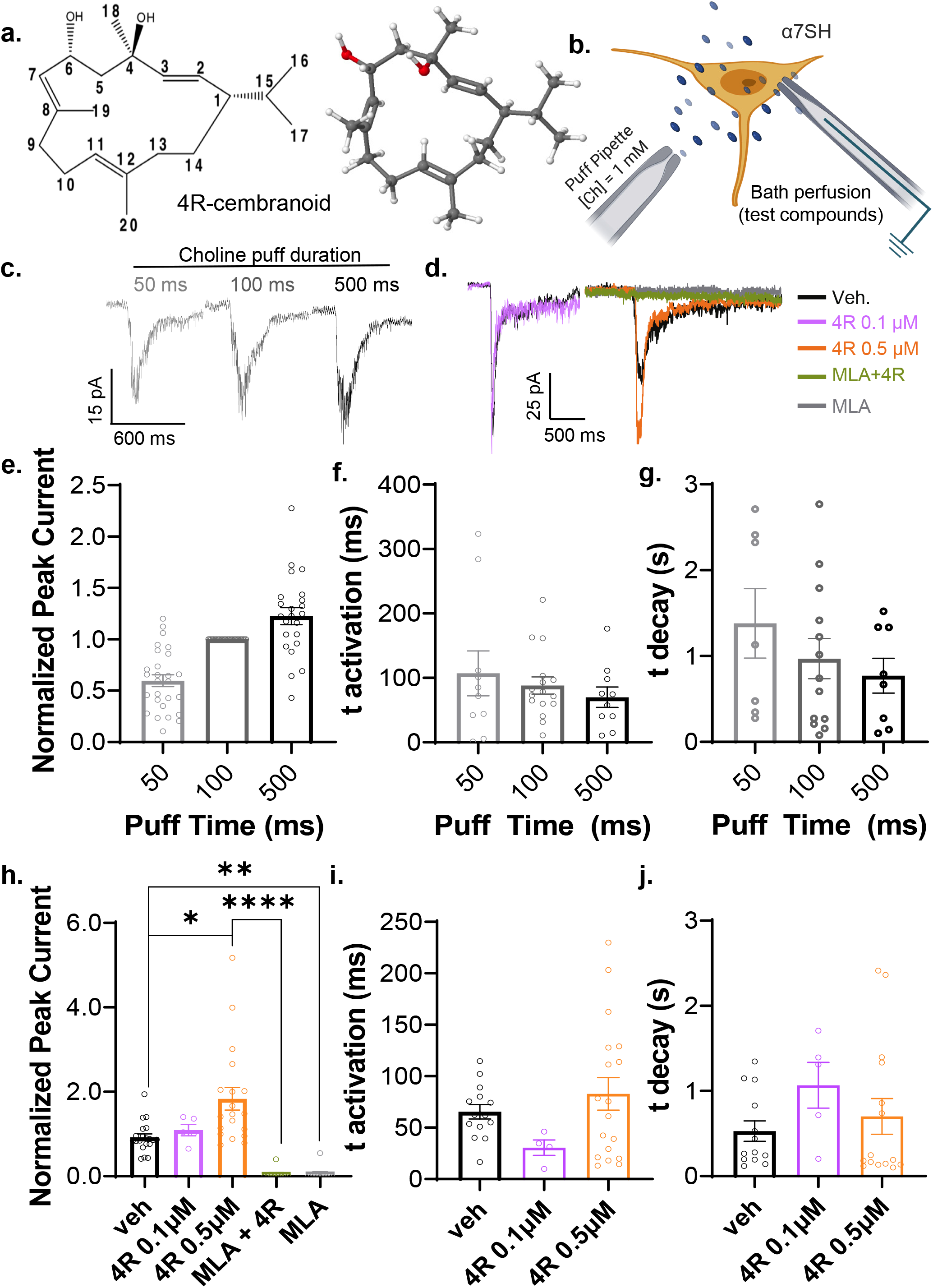
4R potentiates α7 nAChRs-mediated choline-evoked currents. **(a)** Chemical structure (left panel) and 3D molecular model (right panel) of 4R. Oxygen, hydrogen, and carbon atoms are colored red, white, and grey, respectively. **(b)** Schematic of the experimental approach. Responses were evoked by 1 mM choline (Ch) puffs and recorded using whole-cell voltage-clamp in α7SH. **(c)** Representative traces of responses to choline puffs of increasing durations. **(d)** Representative traces of choline-evoked currents to 100 ms puffs after bath application of either vehicle, 4R 0.1 μM, 4R 0.5 μM, MLA 10 μM + 4R 0.5 μM or MLA 10μM. **(e)** dose-dependent increases in peak choline- evoked currents, normalized to 100 ms puff duration. Kruskal-Wallis test; n = 23-27 cells per dose; p < 0.0001 **(f-g)** time of activation **(f)** and deactivation **(g)** of responses to different choline puff durations; n=7-16 cells per dose. **(h-j)** 4R dose response. **(h)** peak choline-evoked currents normalized to pre-drug amplitudes. Kruskal-Wallis test (p < 0.0001) followed by Dunn’s multiple comparison test: veh vs 4R 0.5 μM, p = 0.0389 (*); veh vs MLA 10 μM, p = 0.0027 (**); 4R 0.5 μM vs MLA 10 μM + 4R 0.5 μM, p < 0.0001 (****). (**i-j**) time of activation **(i)** and deactivation (**j**) of choline-evoked currents after bath application of treatments. All values are expressed as mean ± SEM.

## Materials and Methods

### Mice

Male and female C57BL/6J mice (8-17 weeks old; Jackson Labs) were housed in groups of 3-5 littermates with *ad libitum* access to food and water under a 12h/12h light/dark cycle (lights on from 9:00 am to 9:00 pm). One week prior to experiments, mice were transferred in pairs to a new home-cage divided with perforated plexiglass. Mice were handled daily five days before behavioral testing as previously described (Hurst and West 2010). Procedures performed with mice were approved by the Animal Care and Use Committee of the National Institute of Neurologic Disorders and Stroke and the National Institute of Deafness and other Communication Disorders per the National Health Institutes’ (NIH) guidelines. All behavioral experiments were performed by an investigator blind to experimental treatments. Animals were randomly assigned to individual experimental groups. Male and female mice were never tested simultaneously in the same behavior room. All experimental procedures were replicated at least 2 times.

### Inflammatory pain models and behavioral tests

#### CFA paw injections

Mice were lightly anesthetized with isoflurane (0.5%–1% at a flow rate of 0.5 L/min) and 10 μL of Complete Freund’s Adjuvant (CFA, F5881, Sigma Aldrich) was injected subcutaneously into the plantar area of the right hind paw using an insulin syringe (Sure Comfort 29g x 1/2” 3/10cc). Mice with different sexes were evaluated on separate days. Day 0 is defined as the day of CFA injection. Drug solutions used for systemic injections (vehicle or 4R) were made fresh on the day of administration, and a single subcutaneous (s.c.) injection of 56 - 70 μL was administered into the loose skin over the shoulder on day 1. CFA-induced inflammation and hyperalgesia were assessed on day 1 before s.c. injection of vehicle or 4R (1, 6, or 15 mg/kg body weight). The doses of 4R were selected based on previous pharmacokinetic studies in rats (Vélez-Carrasco et al. 2015), extrapolated to mice to account for species differences as previously described (Nair and Jacob 2016). The effects of s.c. treatment with 4R or vehicle on CFA-induced inflammation and hyperalgesia were assessed on days 1, 2, 7 and 8 after the injections. Day 1 effects were measured 2-5 hours after s.c. injections.

For experiments using the α7nAChR-selective antagonist, Methyllycaconitine (MLA), mice received a single vehicle or MLA (10 mgIkg body weight) injection subcutaneously 15 min prior to the injection of vehicle or 4R (15 mg/kg body weight). This dose of MLA has been previously shown to block α7nAChRs-dependent anti-nociception effectively (Freitas et al. 2013, El Nebrisi et al. 2018). The rest of the experiments were performed identically to the 4R experiments described above.

The dorsal-ventral diameter of the CFA-treated and untreated hind paws were measured as an indirect measurement of inflammation using a micro caliper (UX-97152- 17, Cole Palmer). For all behavioral tests, animals were individually placed inside a white plexiglass testing chamber (11×11×13 cm) on an elevated mesh (for acetone and von Frey tests) or a glass platform heated to 30°C (for the heat test).

#### von Frey

Withdrawal thresholds to tactile stimulation, defined as paw withdrawal followed by a brief shake or lick of the paw, were measured using graded monofilaments (North Coast Medical, Inc. San Jose, CA) after animals were habituated to the testing chamber and room for 3 hours, as previously described (Wilson et al. 2019). Starting with the smallest filament, the tip was pressed against the plantar area of the hind paw until it bent at 30° for approximately two seconds. The procedure was repeated five times per filament. The filament that elicited paw withdrawal at least three out of five times was recorded as the mechanical threshold for that trial. The average of three trials was considered the mechanical threshold of the paw tested.

#### Acetone test

Cold sensitivity was assessed by briefly applying an acetone drop to the plantar region of the hind paw and scoring the nocifensive response, as previously described (Wilson et al. 2019). The score system ranges from 0 to 2, indicating the following: 0 = no reaction or an immediate transient lifting or shaking of the hind paw that subsides immediately, 1 = lifting, licking, and/or shaking of the hind paw, which continues beyond the initial application, but subsides within 5 s, and 2 = protracted, repeated lifting, licking, and/or shaking of the hind paw. The response to acetone application was observed for approximately one minute and scored. The average of three stimulations per hind paw represents the score of the paw tested.

#### Hargreaves test

The Hargreaves test was used to assess sensitivity to heat, as previously reported (Wilson et al. 2019). Animals were habituated for one hour in ventilated testing chambers placed on a heated glass surface (30°C). Paw withdrawal latencies were measured after stimulating the hind paw with a heat light aimed at the center of the plantar surface using an active intensity of 25 (intensity of light source as defined by the manufacturer; IITC Life Science, Woodland Hills, CA). Three to five stimulations per hind paw were logged, and the average was reported.

#### Formalin Test

The formalin test was performed in male and female mice as previously described (Tjolsen et al. 1992). Animals were habituated to white plexiglass testing chambers (11×11×13 cm) with clear plexiglass floors for 1 hour. A mirror was placed at a 45° angle under the testing chamber to allow complete visualization of the hind paws. Following habituation, 4R (1, 6, or 15 mg/kg in 43-75 total vol) or vehicle (100% DMSO) was subcutaneously injected into the interscapular area, and the mouse was immediately returned to the testing chamber. Fifteen min after 4R or vehicle administration, animals were injected subcutaneously with 10μL of 2% formalin (Formalin, 252549, Sigma Aldrich) into the plantar surface of the right hind paw. Immediately following intraplantar injection of formalin, time spent in nociceptive behaviors (defined as licking, shaking, and lifting of formalin-injected paw) was recorded in five-minute intervals for 1 hour. To assess the potential anti-inflammatory effects of 4R, the dorsal-ventral diameter of the right hind paw was measured using a micro caliper a day before and immediately following the end of the formalin test. Drug solutions used for systemic (vehicle or 4R) and paw (formalin) injections were made fresh on the day of administration.

### Cell Culture

SH-SY5Y cells overexpressing functional human α7nAChR (α7SH) (Charpantier et al. 2005) were cultured in 1:1 EMEM/Ham’s F12 with NEAA, 2 mM L-glutamine, 15% FBS, 100 μg/mL G418, 100 units/mL Penicillin and 100 μg/mL Streptomycin on 75 cm^3^ tissue culture flasks in an incubator at 37°C and 5% CO_2_. Cells were subsequently seeded into Poly-L-lysine coated glass coverslips at a density of approximately 140 cellslmm^2^ and placed in a 24 well plate as previously described (Shipley et al. 2016). Cells used for electrophysiology were between passages 12 and 20, and recordings were performed 5-10 days after plating.

### Electrophysiology

Choline-evoked currents were recorded from α7SH cells using the whole-cell voltage-clamp technique with a Multiclamp 700B and Digidata 1500 from Molecular Devices. While holding the cells at −70 mV, current traces were acquired at 10 kHz and low pass filtered at 2 kHz using the pClamp 10.7 software. Using a borosilicate glass tube with filament (Warner Instruments), recording pipettes with a resistance of 4-6 MΩ were obtained using the SU-P97 puller (Sutter Instrument Company). Recordings were carried out at room temperature (24-25°C). The external solution consisted of (in mM): NaCl 130, KCl 5, MgCl2 2, CaCl2 2, HEPES 10, and Glucose 25, filtered and pH adjusted to 7.4 with NaOH; theoretical osmolarity: 317 mOsm; real average osmolarity: 308. The internal solution was composed of (in mM): K-gluconate 140, NaCl 5, MgCl2 1.5, CaCl2 0.5, EGTA 5, HEPES 10, Mg-ATP, filtered and pH adjusted to 7.4 with KOH; theoretical osmolarity: 315mOsm; real average osmolarity: 294 mOsm.

All drugs except choline were applied by bath superfusion (3 mL/min) for 90 seconds before recording choline-evoked currents. Choline 1 mM was “puff” applied (15-20 psi; puff durations of 50, 100, 500, and 800 ms) using a pressurized system (Automate Scientific) with a glass pipette with a tip diameter of approximately 20-30 μm, at approximately 150 μm from the cell. The experimental protocol consisted of three different choline (1mM) puff durations (50, 100, 500 ms) applied under different bath treatments (pre-drug, vehicle, or 4R) with an interval of 30 s and 120 s between puff times and treatments, respectively. The order of drug bath perfusions (vehicle or 4R) was counterbalanced to exclude potential time-dependent artifacts.

### Drugs

The tobacco cembranoid (1S,2E,4R,6R,7E,11E)-2,7,11-cembratriene-4,6-diol (4R) was obtained from El Sayed Research Foundation; University of Louisiana–Monroe, College of Pharmacy and prepared as previously described (El Sayed et al. 2008). The purity of the batch used for these experiments was more than 98%, determined by NMR and TLC. 4R was dissolved in DMSO (D2650; Sigma Aldrich) to make a stock solution with a final concentration of 20 mM. The α7nAChR-selective antagonist MLA (1029; Tocris) was dissolved in saline to make a 10 mM stock solution. For electrophysiology experiments, choline (C1879; Sigma Aldrich) was dissolved in external solution to a final concentration of 1 mM and stored at 4°C. Both 4R (20 mM) and MLA (10 mM) stock solutions were diluted in external solution to a final concentration of 0.1, 0.5 or 1.0 μM for 4R and 10 μM for MLA.

For behavioral experiments, a 2.5 mM 4R solution was made from the 4R (20 mM) stock solution to produce the lowest 4R dose injected (1 mg/kg body weight). 4R (20mM) and MLA (10mM) stock solutions were used for the higher doses of 4R (6 and 15 mg/kg body weight) and MLA (10 mg/kg body weight). The concentrations used for 4R electrophysiological experiments were determined via pilot experiments. The concentrations of choline and MLA were based on previous electrophysiological studies demonstrating that these concentrations effectively elicit (or block) choline responses in heterologous systems (Papke et al. 1996, Sokolova et al. 2005, Serres and Carney 2006)

### Data analysis and statistics

Prior to analysis, electrophysiological traces were low-pass filtered offline with an 8-pole RC filter at 1 kHz using ClampFit 10.7 (Molecular Devices). Four distinct parameters were analyzed from each choline-evoked response: peak current, defined as peak amplitude relative to baseline; activation time, defined as rise time from 10% to 90% of the peak amplitude; and decay time, defined as decay time from 90% to 10% of peak amplitude). Peak currents were normalized to responses to 100 ms puff durations within cells for the choline dose-response curve. For 4R and MLA experiments, peak currents were normalized to pre-drug responses within each cell.

All behavioral and electrophysiological data are expressed as mean ± standard error of the mean (S.E.M.). Statistical analyses were performed using Prism (v. 9, GraphPad Software Inc., La Jolla, CA, U.S.A.). For normally distributed data, one-way and two-way ANOVA tests were used for one and two variable studies, respectively, followed by Sidak’s multiple comparison test. Non-normal distributed data were evaluated using the Kruskal-Wallis test followed by Dunnett’s multiple comparison test. P-values < 0.05 were recognized significant.

## Results

### 4R potentiates choline-evoked currents mediated by α7nAChRs

To explore the effects of 4R on α7nAChR-mediated currents, we used a pressurized puff system and whole-cell voltage-clamp recordings of choline-evoked currents on SH-SY5Y overexpressing α7 (α7SH cells) (**Fig. 1b**). As illustrated in **Fig. 1c**, puff application of choline (1 mM) reliably elicited responses with fast onsets (~90 ms) and decays (~1 s) that are stereotypical of α7nAChR-mediated currents at all puff durations. Quantification of peak current amplitudes further revealed that increasing the duration of choline puffs results in dose-dependent increases in choline-evoked currents without affecting the kinetics of the responses (**Fig. 1e-g**).

To determine if 4R modulates choline-evoked currents in α7SH cells, we recorded whole-cell choline-evoked responses to 100 ms puffs of 1mM choline while applying 4R (0.1-0.5 μM) or vehicle in the bath. As illustrated in **Fig. 1d**, bath application of 4R significantly (p < 0.05) increased the peak amplitudes of choline-evoked currents at 0.5 μM but not 0.1 μM 4R, demonstrating that 4R positively modulates choline-evoked responses in α7SH cells in a dose-dependent manner (**Fig. 1h**). The kinetics of activation and decay of choline-evoked currents were unaltered by bath application of 4R compared to vehicle treatment (**Fig. 1i-j**). Importantly, bath application of 4R did not elicit measurable responses in α7SH cells in the absence of choline, demonstrating that it acts as a positive modulator and not an agonist.

The selectivity of choline-evoked responses to α7nAchRs was assessed by pre-treatment with the α7nAChR-selective antagonist MLA (10 nM) in the bath, in the presence or absence of 0.5 μM 4R. As shown in **Fig. 1d and 1h**, choline-evoked responses were blocked by bath application of MLA (10 nM) in α7SH cells treated with 4R 0.5 μM as well as those treated with the vehicle control, demonstrating that α7nAChRs mediate choline-evoked currents in these cells. Together, these results demonstrate that 4R functions as a positive allosteric modulator of α7nAChRs, potentiating choline-evoked currents without altering activation or decay kinetics in α7SH cells.

### Systemic administration of 4R decreases CFA-induced thermal hypersensitivity

Previous studies have shown that α7nAChRs modulate nociceptive responses (Hone and McIntosh 2018), with positive allosteric modulators decreasing injury-induced hypersensitivity (Freitas et al. 2013, El Nebrisi et al. 2018, Nielsen et al. 2020). The next set of experiments aimed at testing the hypothesis that 4R-mediated increases in cholinergic signaling lead to decreases in injury-induced hypersensitivity in male mice. To do this, we used the CFA model of inflammatory pain, which has been shown to cause swelling in the injected paw as well as hypersensitivity to cold, heat, and tactile stimuli (Medhurst et al. 2008, Fehrenbacher et al. 2012). The effects of a one-time systemic administration of 4R on cold, heat, and tactile sensitivity in the CFA-injected and uninjected hind paws were measured at different time-points using acetone, Hargreaves and von Frey test, respectively (**Fig. 2a**). Three different doses of 4R (1, 6, and 15 mg/kg) or vehicle was injected subcutaneously one day after the CFA injection were evaluated.

**Figure 2:**
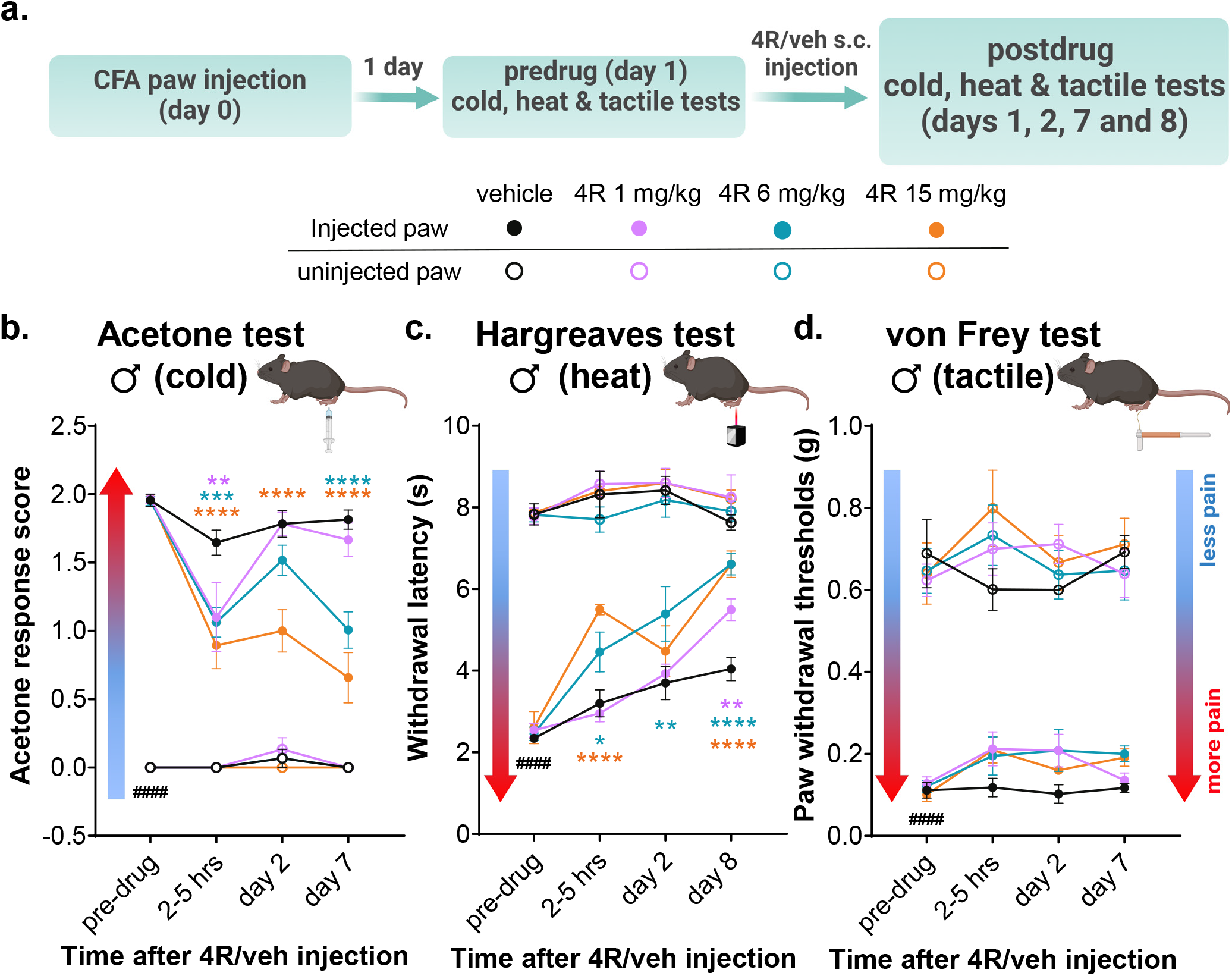
Systemic 4R administration reduces CFA-induced thermal but not tactile hypersensitivity. **(a)** Experimental timeline for acetone (cold), Hargreaves (heat) and von Frey (tactile) tests in CFA-injected and uninjected hind paws before and at different times after 4R (1, 6 and 15 mg/kg) or vehicle administration. **(b)** Acetone response score, **(c)** withdrawal latency to heat stimulation, and **(d)** withdrawal threshold to tactile stimulation. All values are expressed as mean ± SEM. n = 4-9 animals per treatment and test. Two-way ANOVA followed by Dunnett’s multiple comparison test. CFA-injected paw vs uninjected paw in pre-drug vehicle (veh) conditions, p < 0.0001 (####); CFA-injected paws in vehicle vs 4R systemic treatment of the same time-point p < 0.0001 (****), p < 0.01 (**). Results of multiple comparisons between vehicle and 4R 1 mg/kg, 6 mg/kg or 15 mg/kg are shown in purple, teal and orange asterisks, respectively.

Consistent with previous reports (Medhurst et al. 2008, AlSharari et al. 2013), subcutaneous injection of CFA into the hind paw induced hypersensitivity to cold, heat and tactile stimuli in the injected paw, compared to the uninjected hind paw (**Fig. 2b-d**). Thus, prior to 4R treatment (0-hour time-point), the response score to acetone stimulation was significantly (p < 0.0001) higher (**Fig. 2b**) and the withdrawal latencies to heat stimulation were significantly (p < 0.0001) shorter (**Fig. 2c**) in CFA-treated hind paws, compared to the untreated hind paws. Similarly, paw withdrawal thresholds in response to tactile stimulation were significantly (p < 0.0001) lower in the CFA-injected paw compared to the uninjected hind paw (**Fig. 2d**). As expected, CFA-induced hypersensitivity to cold, heat and tactile stimulation was observed in the injected hind paws of control mice (systemically treated with vehicle) for 8 days after CFA injection.

As illustrated in **Fig. 2b-c**, a single injection of 4R reduces CFA-induced hypersensitivity to cold and heat stimuli in a dose and time-dependent manner. Following the administration of the lowest dose of 4R (1 mg/kg), for example, significant (p < 0.0001) decreases in response scores to acetone stimulation were observed in the CFA-injected paw 2-5 hours after 4R treatment, compared to vehicle-treated mice (**Fig. 2b**). The effect of a single injection of 4R (1 mg/kg) in cold hypersensitivity, however, was transient as responses in 4R-treated animals were indistinguishable from control vehicle-treated mice when tested at day 2 and 7 after 4R treatment. In contrast to the transient effects observed after administration of the lowest dose of 4R, the highest dose of 4R tested (15 mg/kg) resulted in long-lasting decreases in CFA-induced cold hypersensitivity, manifested as significantly (p < 0.0001) lower response scores to acetone stimulation of the injected paw, compared to response scores in vehicle-treated mice, in all time-points tested (**Fig. 2b**). Administration of 4R at 6 mg/kg of weight resulted in significant (p < 0.0001) decreases in response scores to acetone stimulation in the CFA-treated hind paw 2-5 hours and on day 7 after the single injection of 4R but not on day 2 after treatment.

Evaluation of CFA-induced hypersensitivity to heat stimuli revealed that 4R also decreases heat hypersensitivity in the inflamed paw in a time and dose-dependent manner (**Fig. 2c**). Mice treated with the lowest 4R dose (1 mg/kg) displayed a significant (p = 0.0075) increase in paw withdrawal latency in response to heat stimulation (less hypersensitivity) of the CFA-injected paw on day 8, but not 2-5 hours or on day 2 after 4R treatment, compared to vehicle-treated animals (**Fig. 2c**). In contrast, administration of the highest dose of 4R (15 mg/kg) showed strong anti-hyperalgesic effects in response to heat stimulation of the injected paw 2-5 hours and day 8 (p < 0.0001), but not on day 2 after 4R treatment. Lastly, animals treated with 4R at 6 mg/kg of body weight displayed significant (p = 0.0036) increases in paw withdrawal latency (less hypersensitivity) on days 2 and 7, but not 2-3 hours post-treatment compared to vehicle-treated animals.

Fig. 2d shows that tactile hypersensitivity, measured as withdrawal thresholds of the CFA-injected paw, was not different from vehicle controls at any dose of 4R and at all times tested after initiation of treatment. In addition, measurement of responses to cold, heat and tactile stimulation of the uninjured hind paw showed that responses in all modalities are indistinguishable in vehicle- and 4R-treated mice throughout the duration of the experiment and independently of the dose tested (**Fig. 2b-d**). Collectively, these results demonstrate that a single systemic administration of 4R decreases inflammation- induced cold and heat hypersensitivity without affecting tactile hypersensitivity or baseline responses in the uninjured paw.

### 4R anti-hyperalgesic effects are mediated via modulation of α7nAChRs

The electrophysiological experiments presented in **Fig. 1h** show that 4R potentiates α7nAChR-mediated currents in α7SH cells. The next set of experiments aimed at testing the hypothesis that 4R-mediated reductions of inflammation-induced behavioral hypersensitivity are mediated via modulation of α7nAChR in-vivo. To do this, the α7nAChRs selective antagonist MLA (10 mg/kg, s.c.) was administered 15 min prior to 4R injection (15 mg/kg, s.c.). CFA-induced cold and heat hypersensitivity was measured on days 7 and 8 after systemic injections, respectively (**Fig. 3a**).

**Figure 3:**
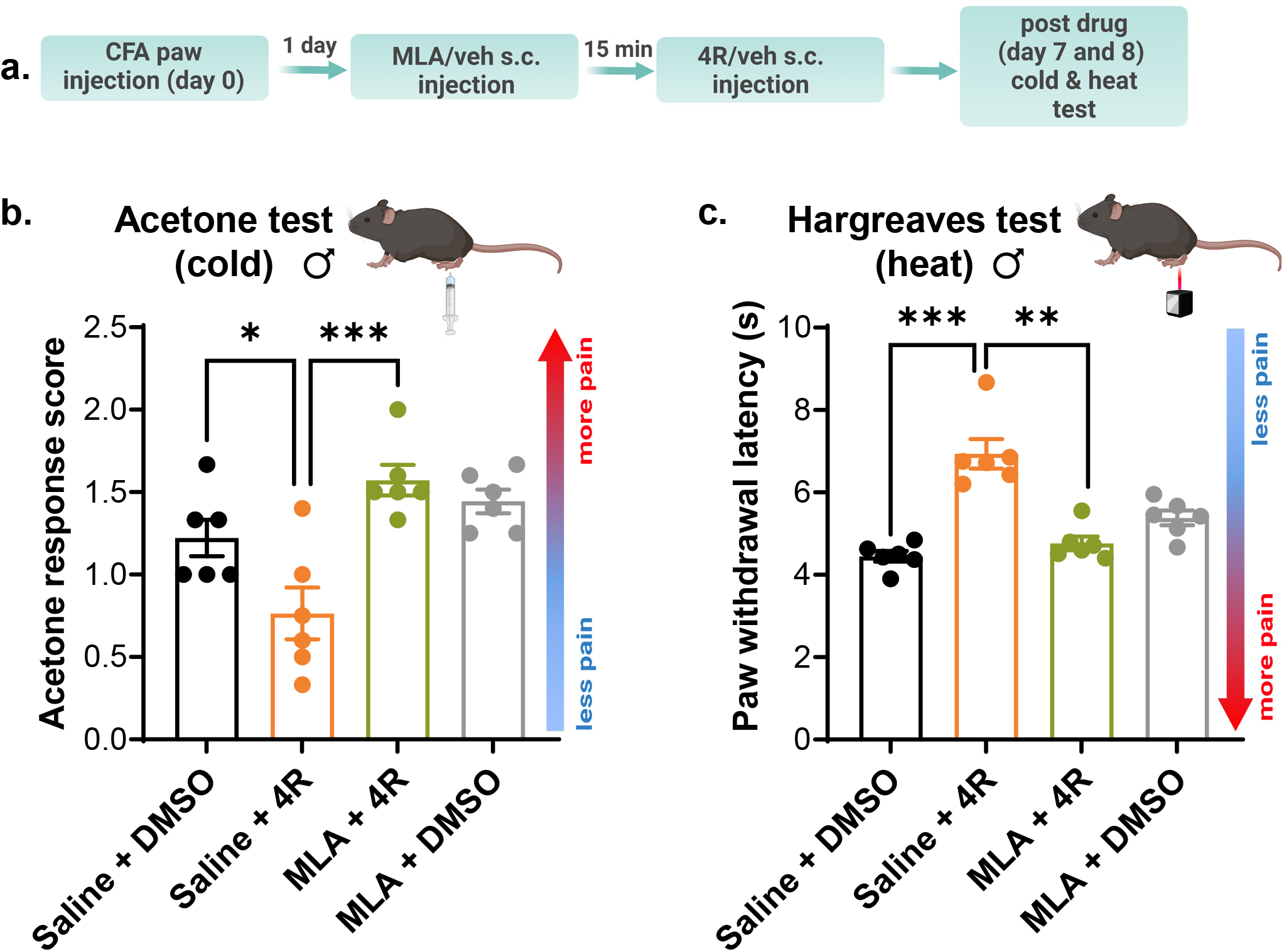
Anti-hyperalgesic effects of 4R are mediated by α7nAChRs. **(a)** Experimental timeline. One day after CFA paw injection, MLA (10 mg/kg) or vehicle (veh) was systemically administered 15 min prior to 4R (15 mg/kg) or veh treatment (s.c.). Acetone (cold) and Hargreaves (heat) tests in CFA-injected and uninjected hind paws were performed 7-8 days after systemic drug administration. **(b)** Acetone response scores; One-way ANOVA (p < 0.001) followed by Šídák’s multiple comparison test: saline + DMSO vs saline + 4R; p < 0.05 (*), saline + 4R vs MLA + 4R, p < 0.001 (***). **(c)** paw withdrawal latencies in response to heat stimulation. Kruskal-Wallis test (p < 0.001) followed by posthoc Dunnett’s multiple comparison test: saline + DMSO vs saline + 4R, p < .001 (***); saline + 4R vs MLA + 4R, p < .01 (**). All values are expressed as mean ± SEM. n = 6 animals per treatment and test.

Consistent with the results presented in **Fig. 2**, animals treated with 4R (15 mg/kg) exhibited lower response scores to acetone stimulation (p = 0.0277) and longer paw withdrawal latencies (p = 0.0002) in response to heat stimulation of the CFA-injected paw compared to vehicle-treated animals, indicating that 4R reduces inflammation-induced cold and heat hypersensitivity **(Fig. 3b-c**). Evaluation of animals pre-treated with the α7nAChRs selective antagonist MLA (10 mg/kg) revealed that MLA pre-treatment prevents the anti-hyperalgesic effects of 4R (**Fig. 3b-c**). Thus, mice systemically treated with MLA prior to 4R show significantly higher response scores to acetone stimulation (p = 0.0002) (**Fig. 3b**) and significantly shorter paw withdrawal latencies in response to heat stimulation (p = 0.0058) (**Fig. 3c**) of the CFA-injected paw compared with responses measured in animals pre-treated with MLA vehicle followed by 4R. Its noteworthy that although MLA abrogates the effect of 4R, MLA per se does not affect the responses to cold or heat stimulation. Together, these results indicate that 4R-induced anti-hyperalgesic effects in response to cold and heat stimuli in a model of inflammatory pain are mediated via modulation of α7nAChRs *in-vivo.*

### 4R reduces pain-related behaviors independently of sex

Several studies report mechanistic sex-related differences in pain modulation, stressing the importance of studying male and female subjects in pre-clinical pain studies (Pabst et al. 2016, Mapplebeck et al. 2017, Rosen et al. 2017, Mogil 2020). In line with this, the next set of experiments aimed to determine whether the 4R anti-hyperalgesic effects described above in males are also observed in females using the same experimental approach (**Fig. 4a**). As illustrated in **Fig. 4b-g**, evaluation of the responses to acetone, Hargreaves and von-Frey tests showed that control female mice display cold, heat and tactile hypersensitivity in CFA-injected paws compared to their respective uninjected paws on days 2 and 7 after systemic vehicle treatment. Thus, response scores to acetone stimulation were significantly (p < 0.0001) higher in CFA-injected hind paws than in the uninjected hind paws (**Fig. 4b, e**). Similarly, paw withdrawal latencies and thresholds to heat and tactile stimulation of the CFA-injected paw were significantly (p < 0.0001) shorter and lower than those measured in the uninjected hind paws (**Fig. 2c, d, f, and g**).

**Figure 4:**
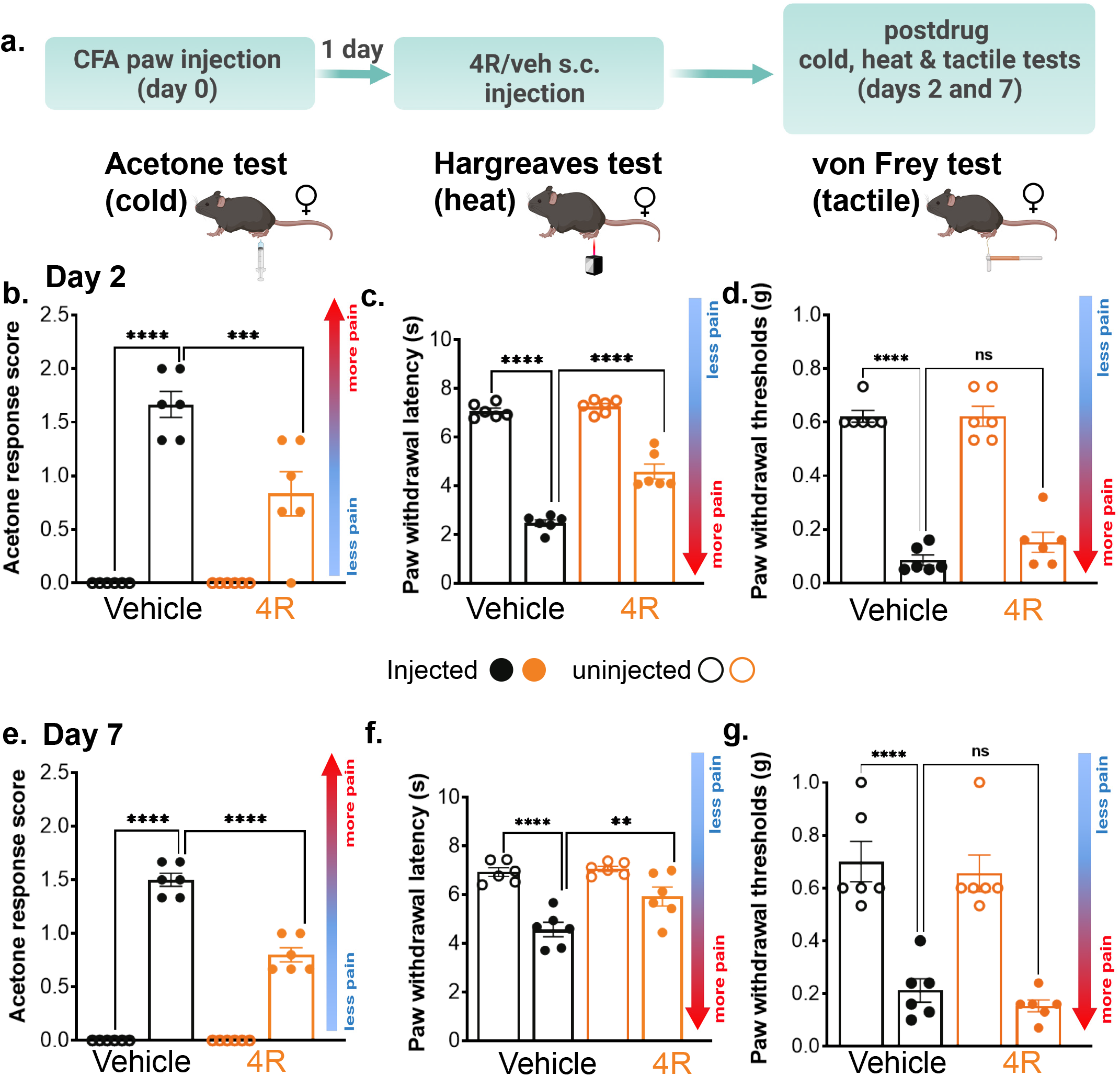
Systemic 4R administration reduces CFA-induced thermal but not tactile hypersensitivity in in female mice. **(a)** Experimental timeline for acetone (cold), Hargreaves (heat) and von Frey (tactile) tests in CFA-injected and uninjected hind paws at days 2 and 7 after 4R (15 mg/kg) or vehicle administration in females. Acetone response score **(b, e)**, withdrawal latency to heat stimulation **(c, f)** and withdrawal threshold to tactile stimulation **(d, g)** on experimental days 2 and 7, respectively. All values are expressed as mean ± SEM. n = 6 animals per treatment and test. One-way ANOVA followed by Šídák’s multiple comparison test; ns = not significant; p < 0.01 (**); p < 0.001 (***); p < 0.0001 (****).

Parallel experiments in females treated with 4R (15 mg/kg) showed that a single s.c. injection of 4R significantly (p < 0.01) decreases CFA-induced cold and heat hypersensitivity compared to mice treated with vehicle on both days 2 and 7 after systemic treatment (**Fig. 4b-e**). Thus, in both days tested, response scores to acetone were significantly (p = 0.0002) lower in the CFA-injected paw of 4R-treated mice than in the injected paws of the vehicle-treated group (**Fig. 4b, c**). Similarly, paw withdrawal latencies to heat stimulation of the injected paw were significantly (p = 0.0056) longer in 4R-treated mice than in those treated with vehicle (**Fig. 4d, e**). In both days tested, sensitivity to cold and heat stimulation of the uninjected hind paw was indistinguishable between vehicle and 4R treated groups.

Consistent with the results observed in males, evaluation of CFA-induced tactile hypersensitivity showed that withdrawal thresholds are comparable in the injected paws of 4R and vehicle-treated mice in both time points evaluated, demonstrating that tactile hypersensitivity is unaltered by 4R treatment (**Fig. 4f, g**). Similarly, paw withdrawal thresholds in the uninjected hind paws are also indistinguishable in 4R and vehicle-treated mice. Altogether, these results demonstrate that a single systemic injection of 4R 15 mg/kg reduces CFA-induced cold and heat hypersensitivity without altering baseline responses or tactile hypersensitivity in both male and female mice.

### Systemic administration of 4R reduces CFA-induced paw edema in male but not female mice

Positive allosteric modulators and agonists of α7nAChRs have been shown to attenuate inflammation-induced paw edema and cytokine release in several rodent models of inflammatory pain (Munro et al. 2012, AlSharari et al. 2013, Freitas et al. 2013, Quadri et al. 2018). In addition, previous studies have shown that systemic treatment with 4R has anti-inflammatory effects in the lipopolysaccharide model of inflammation in mice (Rojas-Colon et al. 2021). To evaluate if the 4R anti-hyperalgesic effects described in the sections above are coupled to decreases in inflammation, we measured paw thickness as an indirect assessment of paw edema and inflammation in CFA-injected and uninjected hind paws before and at different time-points following the systemic administration of various doses of 4R (1, 6 or 15 mg/kg body weight) or vehicle (**Fig. 5a**). As shown in **Fig. 5b-d**, subcutaneous injection of CFA in the hind paw of control male and female mice (systemically treated with vehicle) resulted in significant (p < 0.0001) increases in paw thickness (edema) when compared to their respective uninjected hind paws for the duration of the experiment.

**Figure 5:**
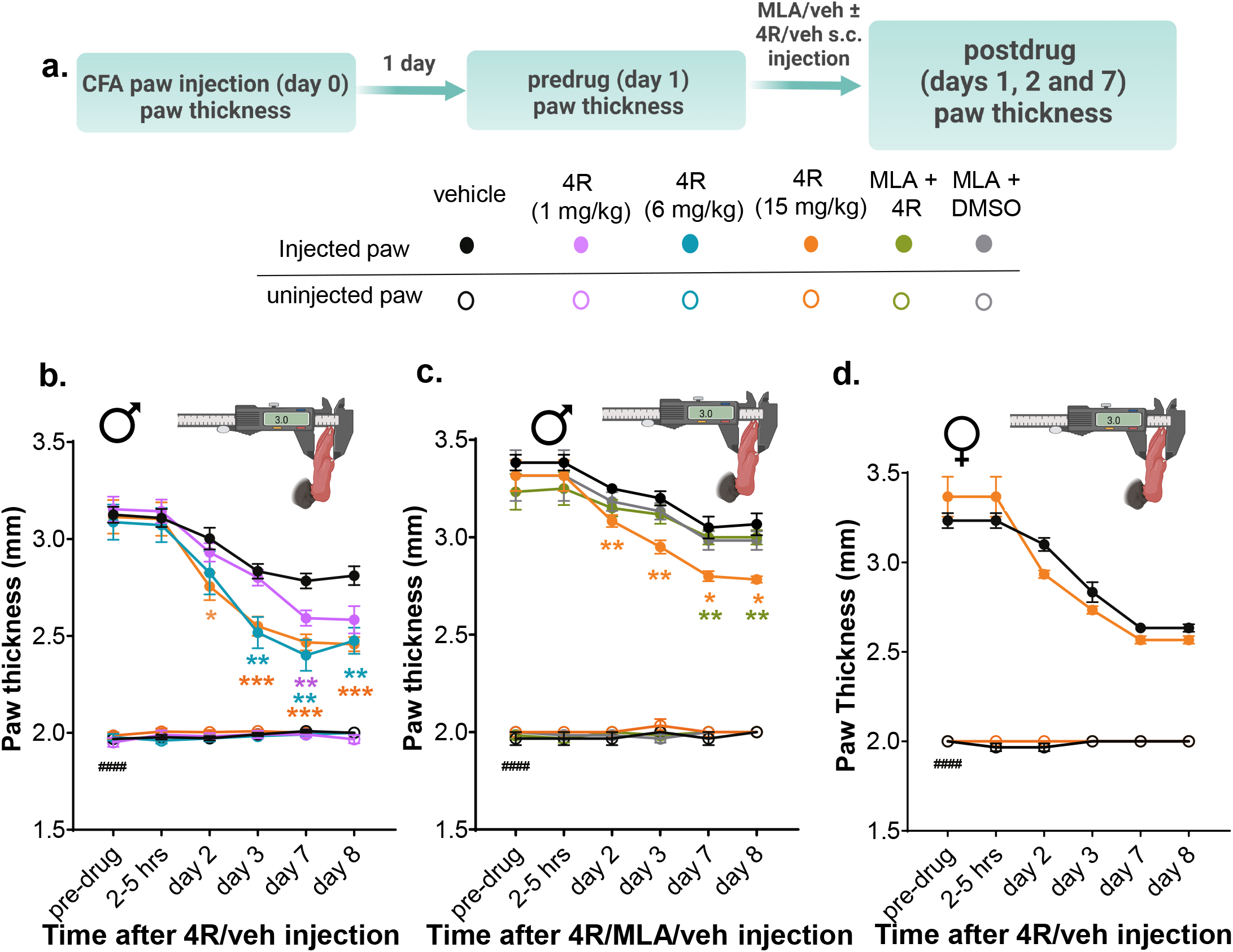
Systemic administration of 4R reduces CFA-induced paw edema in male but not female mice. **(a)** Experimental timeline for CFA-injected and uninjected paw thickness measurements before and at different times after 4R (1, 6 and 15 mg/kg), MLA (10 mg/kg) or vehicle administration in male and female mice. **(b-d)** Thickness measurements of CFA-injected and uninjected hind paws before and 1-8 days after 4R (1, 6 and 15 mg/kg), MLA (10 mg/kg) + 4R (15 mg/kg), MLA (10mg/kg) or vehicle systemic administration. All values are expressed as mean ± SEM. n = 6-16 animals per treatment and test. CFA-injected paw vs uninjected paw in pre-drug vehicle (veh) conditions, p < .0001 (####); CFA-injected paws after treatments of the same time-point p < 0.05 (*), p < .01 (**), p < 0.001 (***), p < .0001 (****). Results of mixed effects analysis followed by Dunnett’s multiple comparison test between vehicle (veh) and 4R 1 mg/kg, 6 mg/kg or 15 mg/kg are shown in purple, teal and orange asterisks, respectively. Results of two-way repeated measures ANOVA followed by Tukey’s multiple comparisons test between veh + 4R (15 mg/kg) and MLA (10 mg/kg) + 4R (15 mg/kg) are shown in green.

Evaluation of paw edema in male mice systemically treated with 4R revealed that a single administration of 4R attenuates CFA-induced paw edema in a dose and time-dependent manner (**Fig. 5b**). Thus, male mice treated with the highest (15 mg/kg) and intermediate (6 mg/kg) doses of 4R showed significant (p < 0.05) reductions in paw thickness compared to vehicle-treated mice. 4R-mediated decreases in CFA-induced paw edema were first observed on day 2 after systemic 4R treatment and lasted for the duration of the experiment (8 days post 4R treatment). In contrast, paw thickness in CFA-injected hind paws of males treated with the lowest dose (1 mg/kg) of 4R was indistinguishable from those measured in vehicle treated mice at all time-points tested. Importantly, paw thickness in uninjected hind paws were comparable in all treatments independently of dose or time-point, demonstrating that 4R-mediated decreases in paw thickness are specific to the inflamed hind paw.

To determine if 4R-mediated reductions in paw edema are via α7nAChR, mice were injected with the α7nAChR selective antagonist MLA (10 mg/kg) 15 min prior to 4R (15 mg/kg) administration. Consistent with the results shown in **Fig 5a**, a single injection of 4R (15 mg/kg) significantly (p < 0.05) reduced CFA-induced paw edema starting at day 2 post 4R treatment when compared to vehicle-injected male mice (**Fig. 5b**). Pre-treatment with MLA (10 mg/kg) prevented the 4R-mediated reductions in paw thickness at days 7 and 8 after treatment. Importantly, CFA-induced paw thickness was not measurably affected by administration of MLA in the absence of 4R compared to vehicle-treated mice, demonstrating that systemic administration of the α7nAChR selective antagonist MLA does not measurably affect CFA-induced paw edema. Collectively, these results show that a single subcutaneous 4R injection reduces CFA-induced paw edema in a dose- and time-dependent manner and that α7nAChRs mediate the observed anti-inflammatory effects.

Parallel experiments were performed in female mice to test if 4R-mediated decreases in inflammation-induced paw edema are sex-specific. Similar to male mice, female mice also developed paw edema after CFA injection that lasted for the duration of the experiment when compared to the uninjected hind paws (p < 0.0001) (**Fig. 5c**). In marked contrast to the results observed in male mice, however, measurement of CFA-induced paw edema in females after systemic treatment with 4R (15 mg/kg) were indistinguishable from those in females treated with vehicle at all time points measured. These results demonstrate that 4R-mediated reductions in inflammation-induced paw edema are specific to males.

### Formalin-induced spontaneous nociceptive responses and paw edema are unaffected by systemic pre-treatment with 4R independently of sex

In the next set of experiments, we used the formalin test as a mouse model of inflammatory pain to evaluate if pre-treatment with a single systemic administration of 4R reduces formalin-induced spontaneous nociceptive behaviors (defined as licking, lifting, and lifting of formalin-injected paw). To do this, we pre-treated mice with a single systemic dose of 4R (1, 6, or 15 mg/kg) or vehicle 15 min before the intraplantar injection of 2% formalin in the hind paw (**Fig. 6a**). Consistent with previous reports (Tjolsen et al. 1992), the time spent in formalin-induced nociceptive behaviors in control vehicle-treated mice was biphasic (**Fig. 6b-c**). Evaluation of the time spent in formalin-induced nociceptive behaviors in male mice pre-treated with different doses of 4R (1, 6, or 15 mg/kg) showed that nociceptive responses are indistinguishable in 4R and vehicle treated animals independently of the dose tested (**Fig. 6b**). Formalin-induced nociceptive responses were also unaffected in female mice systemically pre-treated with 4R (15 mg/kg) compared to vehicle-injected mice (**Fig. 6c**).

**Figure 6:**
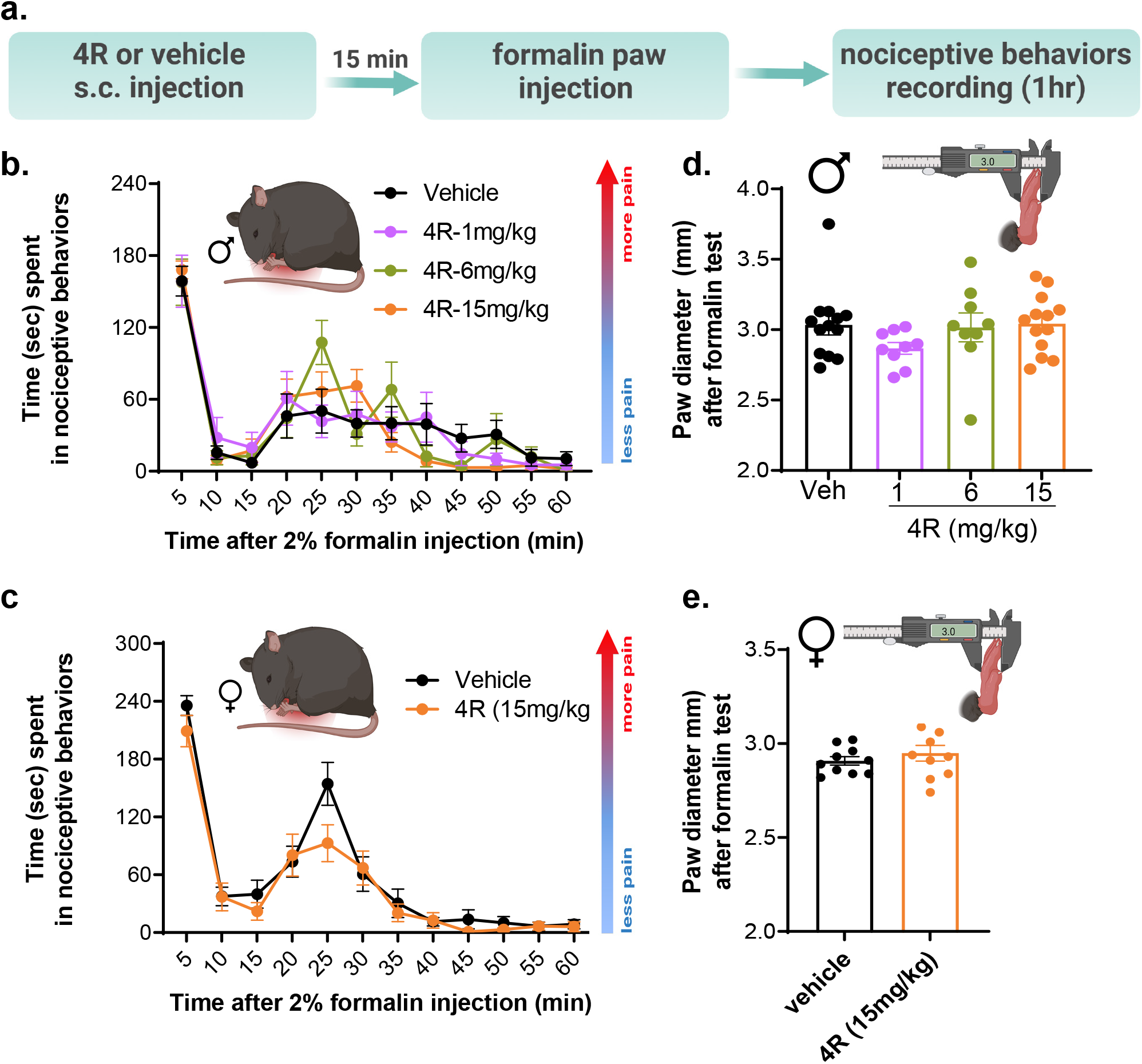
Spontaneous formalin-induced nociceptive responses and paw edema are unaffected by systemic pretreatment with 4R independently of sex. **(a)** Experimental timeline for the formalin test. **(b-c)** Time spent in pain-related behaviors following formalin injection in the right hind paw from 4R (1, 6 or 15 mg/kg) or vehicle-treated male **(b)** and female **(c)** mice. **(d-e)** Formalin-injected paw thickness measurements one day before and immediately after the formalin test in 4R (1, 6 or 15 mg/kg) or vehicle-injected male **(d)** and female **(e)** mice. All values are expressed as mean ± SEM; n= 9-13 males and 7-10 females; data was analyzed using two-way repeated measures ANOVA **(b-c)**, one way ANOVA **(d)** and unpaired t-test **(e)**.

Consistent with the unaltered formalin-induced behavioral responses observed after systemic pre-treatment with 4R (**Fig. 6b-c**), measurements of formalin-induced paw edema showed that paw thickness is also indistinguishable in male and female mice pre-treated with vehicle and 4R independently of the dose tested (**Fig. 6d-e**). Altogether, our results show that formalin-induced spontaneous nociceptive responses and paw edema are unaffected by pre-treatment with 4R in both male and female mice.

## Discussion

Pre-clinical studies have shown that activation of α7nAchRs decreases inflammation and pain-related behaviors in rodents (Wang et al. 2003, Ulloa 2005, Feuerbach et al. 2009, AlSharari et al. 2013, Dineley et al. 2015, Bagdas et al. 2016, Kusuda et al. 2020). Positive allosteric modulators for α7nAchRs that increase endogenous nicotinic transmission without progressive desensitization or loss of receptor function have received increasing attention in recent years as a compelling alternative for pain treatment (Williams et al. 2011). Our results show that 4R potentiates choline-evoked responses *in-vitro* and reduces inflammation-induced thermal hypersensitivity and paw edema *in-vivo,* in a sex-dependent manner, in mice. Together, these findings suggest that 4R might serve as a potential novel therapeutic option to reduce inflammation and pain.

### 4R functions as a positive allosteric modulator of α7nAchRs

Our *in-vitro* electrophysiological experiments show that 4R potentiates choline-evoked α7nAchRs macroscopic currents without altering the kinetics of the response in SH-SY5Y cells overexpressing α7nAChRs (**Fig. 1h-j**). We also demonstrate that 4R does not elicit responses in these cells in the absence of choline and that choline-evoked responses were blocked by selective α7nAchRs antagonist MLA. Together, these results support our hypothesis that 4R functions as a positive allosteric modulator (PAM) of α7nAchRs. PAMs are traditionally classified into Type I and Type II based on their distinctive modulation of α7nAChRs. While both Type I and II PAMs enhance the activity of an agonist, only Type II PAMs decrease receptor desensitization (Hurst et al. 2005, Grønlien et al. 2007, Papke et al. 2009). Based on our results showing that the kinetics of choline-evoked responses are unaltered by bath application of 4R, we predict that 4R functions as Type I PAM. Additional pharmacological and electrophysiological studies to carefully evaluate the effects of 4R on single-channel properties will be needed to determine whether 4R is Type I or Type II PAM.

It is important to note that previous studies have shown that bath application of 25 μM 4R completely inhibits acetylcholine-mediated currents in SH-SY5Y cells overexpressing α7nAChRs (Ferchmin et al. 2013). These results are in marked contrast to our results showing potentiation of choline-evoked responses in these cells when 0.5 μM, a much lower concentration of 4R, was bath applied, with no effect observed when 0.1 μM 4R was used (**Fig. 1h**). Collectively, the results from previous studies and our experiments suggest that 4R-mediated modulation of α7nAChRs is dose-dependent.

### The mechanisms driving 4R-mediated anti-hyperalgesic effects are sexually dimorphic

Our *in-vivo* behavioral experiments using the CFA model of inflammatory pain show that a single systemic administration of 4R reduces inflammation-induced thermal hypersensitivity in both male and female mice (**Figs. 2 and 4**). However, we observed 4R-mediated reductions in CFA-induced paw edema only in males (**Fig. 5**). These results suggest that the mechanisms underlying 4R-mediated anti-hyperalgesic effects are sexually dimorphic.

Previous studies have shown that α7nAchRs are expressed in neuronal and non-neuronal cells (including macrophages) and modulate inflammatory responses and pain-related behaviors (Ulloa 2005, Dineley et al. 2015). Additional studies have further revealed sex-dependent differences in the recruitment of immune cells processing inflammation and pain (Mapplebeck et al. 2017, Rosen et al. 2017, Mogil 2020). Thus, it is possible that the male-specific anti-inflammatory effect of 4R reflects a sexual dimorphism in the types of α7nAchRs-expressing immune cells that modulate inflammatory responses.

Given that an endogenous agonist is required for 4R to potentiate α7nAchRs signaling, an alternative hypothesis is that choline or acetylcholine release in inflammatory states is sexually dimorphic. A third potential source for male-specific anti-inflammatory effects of 4R are sex-dependent differences in intracellular signaling pathways downstream of α7nAchRs activation (Ulloa 2005). While additional experiments are needed to mechanistically address the source of sex-dependent modulation of inflammation by 4R, our results in the present study support the growing body of evidence demonstrating that modulation of inflammatory responses are sexually dimorphic. Our results further stress the importance of including both males and females in pre-clinical pain studies that aim at improving diagnostic and treatment options for pain patients.

We show that 4R reduces inflammation-induced thermal hyperalgesia in females (**Fig. 4**) despite the lack of measurable effects in CFA-induced paw edema (**Fig. 5d**). In males, we observed reductions in CFA-induced thermal hyperalgesia starting at 2.5h post systemic administration of 4R (**Fig. 2b-c**). At this time-point, however, paw edema in males was also measurably unaffected by 4R (**Fig. 5b**). Together, these results suggest that 4R decreases inflammation-induced thermal hyperalgesia via mechanisms that are independent of the mechanisms driving inflammation. In females, the dominant mechanism for 4R anti-hyperalgesic effects seems to be independent of the inflammatory response, as paw edema was unchanged by 4R at all time-points measured. In males, however, 4R might decrease inflammation-induced hyperalgesia independently of the inflammatory response at earlier time-points but might rely on modulation of the inflammatory response at later time-points.

It has been reported that α7nAchRs are expressed in several regions throughout the pain neuraxis and that acute activation of these receptors at different central and peripheral sites modulates pain-related behaviors (Cordero-Erausquin et al. 2004, Naser and Kuner 2018). 4R not only crosses the blood-brain barrier but it seems to be actively transported into the brain (Vélez-Carrasco et al. 2015). We propose that 4R-mediated reduction of inflammation-induced thermal hypersensitivity is mediated via potentiation of neuronal α7nAchRs in both males and females. However, part of the effect of 4R could mediated by other nicotinic receptors or modulated by them (Ferchmin et al. 2005). In males, 4R-mediated decreases in inflammation-induced thermal hypersensitivity are coupled with anti-inflammatory effects, suggesting that 4R might function to potentiate α7nAchRs in non-neuronal cells, including macrophages.

### Prolonged 4R-mediated anti-hyperalgesic and anti-inflammatory effects suggest long-lasting modulation of α7nAchRs downstream signaling pathways

Previous pharmacokinetic and metabolic experiments have shown that the half-life of 4R when administered systemically in rats is between 36 min to 1.5 hours (Velez-Carrasco et al. 2015). These studies further showed no measurable plasma (or brain) levels of 4R 8 hours post administration. Consistent with the reported pharmacokinetics of 4R, our behavioral experiments show 4R-mediated anti-hyperalgesic effects 2.5 hours after a single systemic 4R administration. Notably, our behavioral time-course experiments further showed that 4R anti-hyperalgesic effects are long-lasting, with robust effects measured up to 8 days after systemic treatment with 4R (**Figs. 2 and 4**). Based on previous studies showing that activation of α7nAchRs results in transient calcium influx and subsequent activation of downstream intracellular signaling pathways, we propose that the long-lasting effects of 4R are via modulation of intracellular signaling pathways downstream of α7nAchRs (Ferchmin et al. 2005, Ferchmin et al. 2013).

4R has been previously shown to have neuroprotective effects in various *in-vitro* and *in-vivo* rodent models of neurodegenerative disease and stroke (Ferchmin et al. 2005, Eterović et al. 2011, Ferchmin et al. 2014, Ferchmin et al. 2015, Martins et al. 2015). Whether 4R previously reported neuroprotective functions are also long-lasting as well as the exact duration of the anti-hyperalgesic and anti-inflammatory effects of 4R shown in the present study remains to be determined.

### 4R-mediated anti-hyperalgesia is modality-specific

While inflammation-induced heat and cold hypersensitivity were markedly reduced by a single dose of 4R, tactile hypersensitivity and inflammation-induced spontaneous nociceptive responses were unaffected by 4R treatment independently of dose, time or sex (**Figs. 2, 4 and 6**). These results are consistent with previous studies that show that modulation of pain-related behaviors by activation of α7nAchRs is modality-specific (Dineley et al. 2015).

The experiments in the present study further show that baseline responses to noxious stimulation of the uninjured paw are unaltered by 4R treatment. These results have two important implications. First, it shows that withdrawal responses to heat, cold and tactile stimulation are unaltered by systemic 4R treatment, suggesting that 4R treatment does not elicit overt effects in motor function. Second, it demonstrates that 4R treatment decreases inflammation-induced pathological alterations in behavioral responses without affecting detection and responses to noxious stimuli under physiological conditions, an evolutionarily function essential for survival.

### Concluding Remarks

This study examined the potential use of 4R to decrease inflammation and pain-related hypersensitivity. Our results show that 4R functions as a positive allosteric modulator of α7nAchRs that decreases inflammation and thermal hypersensitivity in a dose-, time- and sex- dependent manner. These results add to the rapidly expanding literature supporting the development and use of pharmacological tools that modulate α7nAchRs to ameliorate pain and inflammation. These results also support previous studies showing that modulation of inflammation is sex-dependent, emphasizing the importance of including males and females in pain studies.

Pain relief by positive allosteric modulators of α7nAChRs has been primarily studied using synthetic compounds. However, several natural or semisynthetic counterparts have also been studied (Ximenis et al. 2021). To the best of our knowledge, 4R is the first positive allosteric modulator isolated from tobacco and cigarette smoke (Saito et al. 1985) that ameliorates pain through potentiation of α7nAChRs. The link between tobacco consumption and pain has been known for decades (Charlton 2004). However, the potential use of tobacco cembranoids for pain treatment has not been evaluated. In addition to 4R, 15 tobacco cembranoids (natural or chemically modified) and analogs with higher potency or higher efficacy have been identified (Eterović et al. 2013). Our results showing that 4R decreases inflammation-induced hyperalgesia for up to 8 days in both male and female mice suggests that tobacco cembranoids offer a promising novel alternative for pain treatment.

### Data Availability

All data in this study is available from the corresponding author.

## AUTHOR CONTRIBUTIONS

Conceptualization, L.G.R.G, V.E., P.F., Y.C.; Investigation, L.G.R., A.M.F.M, T.D.W.; Data Analysis, L.G.R., A.M.F.M, Y.C.; Writing, L.G.R.G, P.F., A.M.F.M., Y.C.; Supervision, Y.C.; Funding Acquisition, Y.C.

## Acknowledgments

This research was supported by the National Center for Complementary and Integrative Health and the National Institute of Neurological Disorders and Stroke Intramural Research Programs. We would like to thank Novartis Pharma AG for kindly donating SH- SY5Y cells overexpressing functional human α7nAChR (α7SH) as well as Dr. Michael Burton for feedback on this study and Dr. Sudhuman Singh for training and assistance with the behavioral experiments. Cartoons in figures were created with BioRender.com.

## References

AlSharari, S. D., K. Freitas and M. I. Damaj (2013). “Functional role of alpha7 nicotinic receptor in chronic neuropathic and inflammatory pain: Studies in transgenic mice.” Biochemical Pharmacology 86(8): 1201–1207.

Bagdas, D., J. L. Wilkerson, A. Kulkarni, W. Toma, S. AlSharari, Z. Gul, A. H. Lichtman, R. L. Papke, G. A. Thakur and M. I. Damaj (2016). “The α7 nicotinic receptor dual allosteric agonist and positive allosteric modulator GAT107 reverses nociception in mouse models of inflammatory and neuropathic pain.” British Journal of Pharmacology 173(16): 2506–2520.

Bektas, N., D. Nemutlu, M. Cam, Y. Okcay, H. Eken and R. Arslan (2020). “The nicotinic modulation of pain.” Pakistan Journal of Pharmaceutical Sciences 33(1): 229–239.

Bektas, N., D. Nemutlu, M. Cam, Y. Okcay, H. Eken and R. Arslan (2020). “Review: The nicotinic modulation of pain.” Pak J Pharm Sci 33(1): 229–239.

Charlton, A. (2004). “Medicinal uses of tobacco in history.” J R Soc Med 97(6): 292–296.

Charpantier, E., A. Wiesner, K.-H. Huh, R. Ogier, J.-C. Hoda, G. Allaman, M. Raggenbass, D. Feuerbach, D. Bertrand and C. Fuhrer (2005). “Alpha7 neuronal nicotinic acetylcholine receptors are negatively regulated by tyrosine phosphorylation and Src-family kinases.” The Journal of neuroscience : the official journal of the Society for Neuroscience 25(43): 9836–9849.

Cordero-Erausquin, M., S. Pons, P. Faure and J.-P. Changeux (2004). “Nicotine differentially activates inhibitory and excitatory neurons in the dorsal spinal cord.” Pain 109(3): 308–318.

Damaj, M. I., E. M. Meyer and B. R. Martin (2000). “The antinociceptive effects of α7 nicotinic agonists in an acute pain model.” Neuropharmacology 39(13): 2785–2791.

Dineley, K. T., A. A. Pandya and J. L. Yakel (2015). “Nicotinic ACh receptors as therapeutic targets in CNS disorders.” Trends Pharmacol Sci 36(2): 96–108.

Dineley, K. T., A. A. Pandya and J. L. Yakel (2015). “Nicotinic ACh receptors as therapeutic targets in CNS disorders.” Trends in pharmacological sciences 36(2): 96–108.

El Nebrisi, E. G., D. Bagdas, W. Toma, H. Al Samri, A. Brodzik, Y. Alkhlaif, K.-H. S. Yang, F. C. Howarth, I. M. Damaj and M. Oz (2018). “Curcumin Acts as a Positive Allosteric Modulator of α 7-Nicotinic Acetylcholine Receptors and Reverses Nociception in Mouse Models of Inflammatory Pain.” Journal of Pharmacology and Experimental Therapeutics 365(1): 190–200.

El Sayed, K. A., S. Laphookhieo, M. Yousaf, J. A. Prestridge, A. B. Shirode, V. B. Wali and P. W. Sylvester (2008). “Semisynthetic and biotransformation studies of (1S,2E,4S,6R,7E,11 E)-2,7,11-cembratriene-4,6-diol.” Journal of Natural Products 71(1): 117–122.

Eterović, V. A., A. Del Valle-Rodriguez, D. Pérez, M. Carrasco, M. A. Khanfar, K. A. El Sayed and P. A. Ferchmin (2013). “Protective activity of (1S,2E,4R,6R,7E,11E)-2,7,11-cembratriene-4,6-diol analogues against diisopropylfluorophosphate neurotoxicity: Preliminary structure–activity relationship and pharmacophore modeling.” Bioorganic & Medicinal Chemistry 21(15): 4678–4686.

Eterović, V. A., D. Pérez, A. H. Martins, B. L. Cuadrado, M. Carrasco and P. A. A. Ferchmin (2011). “A cembranoid protects acute hippocampal slices against paraoxon neurotoxicity.” Toxicology in Vitro 25(7): 1468–1474.

Fehrenbacher, J. C., M. R. Vasko and D. B. Duarte (2012). “Models of Inflammation: Carrageenan-or Complete Freund’s Adjuvant (CFA)-Induced Edema and Hypersensitivity in the Rat.” Current Protocols in Pharmacology 56(1): 5.4.1–5.4.4.

Ferchmin, P., R. Lukas, R. Hann, J. Fryer, J. Eaton, O. Pagan, A. Rodriguez, Y. Nicolau, M. Rosado and S. Cortes (2001). “Tobacco cembranoids block behavioral sensitization to nicotine and inhibit neuronal acetylcholine receptor function.” Journal of neuroscience research 64(1): 18–25.

Ferchmin, P. A., M. Andino, R. Reyes Salaman, J. Alves, J. Velez-Roman, B. Cuadrado, M. Carrasco, W. Torres-Rivera, A. Segarra, A. H. Martins, J. E. Lee, V. A. Eterovic, J. V.-r. Brenda, C. Marime, C. Wilmarie, T.-r. A. Segarra, H. Martins, J. Eun and L. Vesna (2014). “4R-cembranoid protects against diisopropylfluorophosphate-mediated neurodegeneration.” NeuroToxicology 44: 80–90.

Ferchmin, P. A., J. Hao, D. Perez, M. Penzo, H. M. Maldonado, M. T. Gonzalez, A. D. Rodriguez and J. de Vellis (2005). “Tobacco cembranoids protect the function of acute hippocampal slices against NMDA by a mechanism mediated by α4β2 nicotinic receptors.” Journal of Neuroscience Research 82(5): 631–641.

Ferchmin, P. A., O. R. Pagan, H. Ulrich, A. C. Szeto, R. M. Hann and V. A. Eterovic (2009). “Actions of octocoral and tobacco cembranoids on nicotinic receptors.” Toxicon 54(8): 1174–1182.

Ferchmin, P. A., D. Perez, W. Castro Alvarez, M. A. Penzo, H. M. Maldonado and V. A. Eterovic (2013). “gamma-Aminobutyric acid type A receptor inhibition triggers a nicotinic neuroprotective mechanism.” J Neurosci Res 91(3): 416–425.

Ferchmin, P. A., D. Pérez, B. L. Cuadrado, M. Carrasco, A. H. Martins and V. A. Eterović (2015). “Neuroprotection Against Diisopropylfluorophosphate in Acute Hippocampal Slices.” Neurochemical Research 40(10): 2143–2151.

Feuerbach, D., K. Lingenhoehl, H. R. Olpe, A. Vassout, C. Gentsch, F. Chaperon, J. Nozulak, A. Enz, G. Bilbe, K. McAllister and D. Hoyer (2009). “The selective nicotinic acetylcholine receptor α7 agonist JN403 is active in animal models of cognition, sensory gating, epilepsy and pain.” Neuropharmacology 56(1): 254–263.

Freitas, K., F. I. Carroll and M. I. Damaj (2013). “The Antinociceptive Effects of Nicotinic Receptors 7-Positive Allosteric Modulators in Murine Acute and Tonic Pain Models.” Journal of Pharmacology and Experimental Therapeutics 344(1): 264–275.

Freitas, K., S. S. S. Negus, F. I. Carroll and M. I. Damaj (2013). “In vivo pharmacological interactions between a type II positive allosteric modulator of α7 nicotinic ACh receptors and nicotinic agonists in a murine tonic pain model.” British Journal of Pharmacology 169(3): 567–579.

Grønlien, J. H., M. Håkerud, H. Ween, K. Thorin-Hagene, C. A. Briggs, M. Gopalakrishnan and J. Malysz (2007). “Distinct profiles of α7 nAChR positive allosteric modulation revealed by structurally diverse chemotypes.” Molecular pharmacology 72(3): 715–724.

Gurun, M. S., R. Parker, J. C. Eisenach and M. Vincler (2009). “The Effect of Peripherally Administered CDP-Choline in an Acute Inflammatory Pain Model: The Role of α7 Nicotinic Acetylcholine Receptor.” Anesthesia & Analgesia 108(5): 1680–1687.

Hone, A. J. and J. M. McIntosh (2018). “Nicotinic acetylcholine receptors in neuropathic and inflammatory pain.” FEBS Lett 592(7): 1045–1062.

Hone, A. J. and J. M. McIntosh (2018). “Nicotinic acetylcholine receptors in neuropathic and inflammatory pain.” FEBS Letters 592(7): 1045–1062.

Hurst, J. L. and R. S. West (2010). “Taming anxiety in laboratory mice.” Nature Methods 7(10).

Hurst, R. S., M. Hajós, M. Raggenbass, T. M. Wall, N. R. Higdon, J. A. Lawson, K. L. Rutherford-Root, M. B. Berkenpas, W. E. Hoffmann, D. W. Piotrowski, V. E. Groppi, G. Allaman, R. Ogier, S. Bertrand, D. Bertrand and S. P. Arneric (2005). “A novel positive allosteric modulator of the alpha7 neuronal nicotinic acetylcholine receptor: in vitro and in vivo characterization.” The Journal of neuroscience : the official journal of the Society for Neuroscience 25(17): 4396–4405.

Kusuda, R., E. U. Carreira, L. Ulloa, F. Q. Cunha, A. Kanashiro and T. M. Cunha (2020). “Choline attenuates inflammatory hyperalgesia activating nitric oxide/cGMP/ATP-sensitive potassium channels pathway.” Brain Research 1727(November 2018): 146567–146567.

Mapplebeck, J. C. S., S. Beggs and M. W. Salter (2017). “Molecules in pain and sex: a developing story.” Molecular brain 10(1): 9–9.

Martins, A. H., J. Hu, Z. Xu, C. Mu, P. Alvarez, B. D. Ford, K. El Sayed, V. A. Eterovic, P. A. Ferchmin and J. Hao (2015). “Neuroprotective activity of (1S,2E,4R,6R,-7E,11E)-2,7,11-cembratriene-4,6-diol (4R) in vitro and in vivo in rodent models of brain ischemia.” Neuroscience 291: 250–259.

Medhurst, S. J., J. P. Hatcher, C. J. Hille, S. Bingham, N. M. Clayton, A. Billinton and I. P. Chessell (2008). “Activation of the α7-Nicotinic Acetylcholine Receptor Reverses Complete Freund Adjuvant-Induced Mechanical Hyperalgesia in the Rat Via a Central Site of Action.” Journal of Pain 9(7): 580–587.

Mogil, J. S. (2020). “Qualitative sex differences in pain processing: emerging evidence of a biased literature.” Nature Reviews Neuroscience 21(7): 353–365.

Munro, G., R. R. Hansen, H. K. Erichsen, D. B. Timmermann, J. K. Christensen and H. H. Hansen (2012). “The α7 nicotinic ACh receptor agonist compound B and positive allosteric modulator PNU-120596 both alleviate inflammatory hyperalgesia and cytokine release in the rat.” British Journal of Pharmacology 167(2): 421–435.

Nair, A. and S. Jacob (2016). “A simple practice guide for dose conversion between animals and human.” Journal of Basic and Clinical Pharmacy 7(2): 27–27.

Naser, P. V. and R. Kuner (2018). “Molecular, Cellular and Circuit Basis of Cholinergic Modulation of Pain.” Neuroscience 387: 135–148.

Nielsen, B. E., S. Stabile, C. Vitale and C. Bouzat (2020). “Design, Synthesis, and Functional Evaluation of a Novel Series of Phosphonate-Functionalized 1,2,3-Triazoles as Positive Allosteric Modulators of α7 Nicotinic Acetylcholine Receptors.” ACS Chemical Neuroscience 11(17): 2688–2704.

Pabst, M., O. Braganza, H. Dannenberg, W. Hu, L. Pothmann, J. Rosen, I. Mody, K. van Loo, K. Deisseroth, Albert J. Becker, S. Schoch and H. Beck (2016). “Astrocyte Intermediaries of Septal Cholinergic Modulation in the Hippocampus.” Neuron: 1–13.

Papke, R. L., M. Bencherif and P. Lippiello (1996). “An evaluation of neuronal nicotinic acetylcholine receptor activation by quaternary nitrogen compounds indicates that choline is selective for the α7 subtype.” Neuroscience letters 213(3): 201–204.

Papke, R. L., W. R. Kem, F. Soti, G. Y. López-Hernández and N. A. Horenstein (2009). “Activation and desensitization of nicotinic alpha7-type acetylcholine receptors by benzylidene anabaseines and nicotine.” J Pharmacol Exp Ther 329(2): 791–807.

Quadri, M., D. Bagdas, W. Toma, C. Stokes, N. A. Horenstein, M. I. Damaj and R. L. Papke (2018). “The Antinociceptive and Anti-Inflammatory Properties of the α 7 nAChR Weak Partial Agonist p-CF 3 N, N-diethyl-N ‘-phenylpiperazine.” Journal of Pharmacology and Experimental Therapeutics 367(2): 203–214.

Quick, M. W. and R. A. J. Lester (2002). “Desensitization of neuronal nicotinic receptors.” Journal of Neurobiology 53(4): 457–478.

Rojas-Colon, L. A., P. K. Dash, F. A. Morales-Vias, M. Lebron-Davila, P. A. Ferchmin, J. B. Redell, G. Maldonado-Martinez and W. I. Velez-Torres (2021). “4R-cembranoid confers neuroprotection against LPS-induced hippocampal inflammation in mice.” J Neuroinflammation 18(1): 95.

Rosen, S. F., B. Ham, S. Drouin, N. Boachie, A. J. Chabot-Dore, J. S. Austin, L. Diatchenko and J. S. Mogil (2017). “T-cell mediation of pregnancy analgesia affecting chronic pain in mice.” Journal of Neuroscience 37(41): 9819–9827.

Rowley, T. J., A. McKinstry, E. Greenidge, W. Smith and P. Flood (2010). “Antinociceptive and anti-inflammatory effects of choline in a mouse model of postoperative pain.” British Journal of Anaesthesia 105(2): 201–207.

Saito, Y., H. Takizawa, S. Konishi, D. Yoshida and S. Mizusaki (1985). “Identification of cembratriene-4,6-diol as antitumor-promoting agent from cigarette smoke condensate.” Carcinogenesis 6(8): 1189–1194.

Schappert, S. M. and C. W. Burt (2006). “Ambulatory care visits to physician offices, hospital outpatient departments, and emergency departments: United States, 2001-02.” Vital Health Stat 13(159): 1–66.

Serres, F. and S. L. Carney (2006). “Nicotine regulates SH-SY5Y neuroblastoma cell proliferation through the release of brain-derived neurotrophic factor.” Brain research 1101(1): 36–42.

Shipley, M. M., C. A. Mangold and M. L. Szpara (2016). “Differentiation of the SH-SY5Y Human Neuroblastoma Cell Line.” Journal of Visualized Experiments (108): 1–11.

Sokolova, E., C. Matteoni and A. Nistri (2005). “Desensitization of neuronal nicotinic receptors of human neuroblastoma SH-SY5Y cells during short or long exposure to nicotine.” British Journal of Pharmacology 146(8): 1087–1095.

Targowska-Duda, K. M., D. Feuerbach, G. Biala, K. Jozwiak and H. R. Arias (2014). “Antidepressant activity in mice elicited by 3-furan-2-yl-N-p-tolyl-acrylamide, a positive allosteric modulator of the α7 nicotinic acetylcholine receptor.” Neuroscience Letters 569: 126–130.

Tjolsen, A., O. G. Berge, S. Hunskaar, J. H. Rosland and K. Hole (1992). “The formalin test: an evaluation of the method.” Pain 51(1): 5–17.

Ulloa, L. (2005). “The vagus nerve and the nicotinic anti-inflammatory pathway.” Nat Rev Drug Discov 4(8): 673–684.

Ulloa, L. (2005). “The vagus nerve and the nicotinic anti-inflammatory pathway.” Nature Reviews Drug Discovery 4(8): 673–684.

Uteshev, V. V. (2014). “The therapeutic promise of positive allosteric modulation of nicotinic receptors.” European Journal of Pharmacology 727: 181–185.

Velez-Carrasco, W., C. E. Green, P. Catz, A. Furimsky, K. O’Loughlin, V. A. Eterovic and P. A. Ferchmin (2015). “Pharmacokinetics and Metabolism of 4R-Cembranoid.” PLoS One 10(3): e0121540.

Vélez-Carrasco, W., C. E. Green, P. Catz, A. Furimsky, K. O’Loughlin, V. A. Eterović and P. A. Ferchmin (2015). “Pharmacokinetics and Metabolism of 4R-Cembranoid.” PLOS ONE 10(3): e0121540–e0121540.

Wang, H., M. Yu, M. Ochani, C. A. Amelia, M. Tanovic, S. Susarla, J. H. Li, H. Wang, N. Yang, L. Ulloa, Y. Al-Abed, C. J. Czura and K. J. Tracey (2003). “Nicotinic acetylcholine receptor α7 subunit is an essential regulator of inflammation.” Nature 421(6921): 384–388.

Williams, D. K., J. Wang and R. L. Papke (2011). “Positive allosteric modulators as an approach to nicotinic acetylcholine receptor-targeted therapeutics: Advantages and limitations.” Biochemical Pharmacology 82(8): 915–930.

Wilson, T. D. T. D., S. Valdivia, A. Khan, H.-S. H. S. Ahn, A. P. A. P. Adke, S. M. Gonzalez, Y. K. Y. K. Sugimura, Y. Carrasquillo and S. Martinez Gonzalez (2019). “Dual and Opposing Functions of the Central Amygdala in the Modulation of Pain.” Cell Reports 29(2): 332–346.e335.

Ximenis, M., J. Mulet, S. Sala, F. Sala, M. Criado, R. González-Muñiz and M. J. P. de Vega (2021). “Natural polyhydroxy flavonoids, curcuminoids, and synthetic curcumin analogs as α7 nAChRs positive allosteric modulators.” International Journal of Molecular Sciences 22(2): 1–17.

